# PIP_2_-Dependent Thermoring Basis for Cold-Sensing of the TRPM8 Biothermometer

**DOI:** 10.1101/2022.01.02.474742

**Authors:** Guangyu Wang

## Abstract

The menthol sensor TRPM8 can be activated by cold and thus serves as a biothermometer in a primary afferent sensory neuron for innocuous-to-noxious cold detection. However, the precise structural origins of specific temperature thresholds and sensitivity have remained elusive. Here, a grid thermodynamic model was employed to examine if the temperature-dependent noncovalent interactions found in the 3D structures of thermo-gated TRPM8 could assemble into a well-organized fluidic grid-like mesh network, featuring the constrained grids as the thermorings for cold-sensing in response to PIP_2_, Ca^2+^ and chemical agents. The results showed that the different interactions of TRPM8 with PIP_2_ during the thermal incubation induced the formation of the biggest grids with distinct melting temperature threshold ranges. Further, the overlapped threshold ranges between open and pre-open closed states were required for initial cold activation with the matched thermo-sensitivity and the decrease in the systematic thermal instability. Finally, the intact anchor grid near the lower gate was important for channel opening with the active selectivity filter. Thus, PIP_2_-dependent thermorings in TRPM8 may play a pivotal role in cold sensing. (176 words)

## 1. INTRODUCTION

Transient receptor potential (TRP) melastatin-8 (TRPM8) is a Ca^2+^-permeable ion channel. Its activation follows Boltzmann-type relation, and its voltage activation range is dynamically regulated by temperature changes. It serves as a polymodal sensor in response to both physical and chemical stimuli. The physical stimuli include membrane depolarization and cold while the chemical stimuli cover the membrane phosphatidylinositol-4, 5-biphosphate (PIP_2_), lysophospholipids (LPLs) and the cooling agents such as menthol and WS-12 and icilin. ^1-11^ When elevated temperature decreases the menthol-evoked activity of TRPM8, or menthol increases the temperature threshold and the maximal activity temperature of rat TRPM8 (rTRPM8) from 22-26 °C and 8.2 °C to 30-32 °C and 15.6 °C, respectively, the same gating pathway seems to be shared. ^1-2, 12-18^ However, some menthol-sensitive residues such as Y745 and Y1005 in mouse TRPM8 (mTRPM8) are insensitive to cold, challenging the same gating pathway. ^19, 20^ On the other hand, the cold activation of TRPM8 exhibits a high temperature sensitivity Q_10_ of 24-35 (the ratio of rates or open probabilities (P_o_) of an ion channel measured 10 °C apart) at the physiological condition.^12, 21^ Although the cryo-electron microscopy (cryo-EM) structures of closed and open mTRPM8 at 20 °C are available in the absence or presence of Ca^2+^, PIP_2_, and chemical agents such as cryosim-3 (C3) and allyl isothiocyanate (AITC), ^22^ the precise structural factor or motif for the specific PIP_2_-dependent cold temperature threshold and sensitivity is still missing.

Recent studies have demonstrated that a biological macromolecule has a thermoring structure to govern its thermal stability and activity. The thermoring can be a DNA hairpin in a single polynucleotide chain or a 3D topological grid in a systematic fluidic grid-like noncovalent interaction mesh network of a single polypeptide chain. ^23-28^ Such a biochemical or biophysical network can be characterized by a graph theory-based grid thermodynamic model. First, a melting temperature threshold (T_m_) for thermal unfolding of the thermoring from the biggest grid to the smallest one is determined by its strength and size. Second, the grid-based systematic thermal instability (T_i_) at a specific temperature is governed by the ratio of the total grid size to the total noncovalent interactions. Third, a structural thermo-sensitivity (Ω_10_) is controlled by a change in the total grid sizes upon an alteration in the total noncovalent interactions along the same polypeptide chain within a 10 °C interval. Once three calculated parameters match the experimental values including the functional thermo-sensitivity Q_10_, the specific thermoring can be identified as the structural motif or factor for the structural thermostability or the functional thermoactivity. ^24-28^ Following a success in deciphering the lipid-dependent heat response and sensitivity of the TRP vanilloid 1 or 3 (TRPV1 or TRPV3) channel, ^27-28^ it is attractive to use this graphical method to examine such a hypothesis that the temperature-dependent systematic fluidic grid-like noncovalent interaction mesh networks, once identified in the 3D structures of the TRPM8 channel, can be constrained as a thermoring structure from the biggest grid to the smallest one to control the PIP_2_-dependent cold-sensing with specific temperature threshold and sensitivity.

In this computational study, three PIP_2_-dependent biggest grids were first identified near the outer pore loop of mTRPM8. Their melting temperature threshold ranges decreased upon the in turn binding of the chemical agents C3 and AITC together with Ca^2+^. Once the threshold matched the incubation temperature 20 °C, the channel was open with the intact anchor grid near the lower S6 gate to secure the active selectivity filter, the decrease in the systematic thermal instability, and the matched structural and functional thermo-sensitivities. Taken as a whole, although TRPM8 can be finally opened by chemical agents in the presence of Ca^2+^ and PIP_2_, the PIP_2_-dependent thermorings may still serve as the critical structural motif for cold-evoked activation of TRPM8 with the specific temperature threshold and sensitivity.

## 2. MATERIALS AND METHODS

### 2.1 Data mining resources

In this study, cryo-EM structural data of mTRPM8 in three gating states were exploited to prepare the systemic fluidic grid-like noncovalently interacting mesh networks. They included the closed states at 20 °C with PIP_2_ bound (PDB ID, 8E4N, model resolution = 3.07 Å) and with PIP_2_, Ca^2+^, and C3 bound (PDB ID, 8E4M, model resolution = 3.44 Å), and the open state at 20 °C with PIP_2_, Ca^2+^, C3 and AITC bound (PDB ID, 8E4L, model resolution = 3.32 Å). ^22^ UCSF Chimera was used for the thermoring analyses.

### 2.2 The definition of the necessary gating pathway

PIP_2_ is required for TRPM8 opening. ^5-6, 9, 22^ The cryo-EM structure indicates that S4b, TRP domain, and pre-S1 of mTRPM8 form the active PIP_2_ binding pocket. ^22^ The primary structural analysis showed that mTRPM8 with PIP_2_ bound at 20 °C (PDB ID, 8E4N) had F1013 in the C-terminal domain to form a π−π interaction with F744 on S1. On the other hand, Q675 in the pre-S1 domain H-bonded with R998 in the TRP domain in both closed and open states.^22^ Hence, the segment from Q675 in the pre-S1 domain to F1013 in the distal C-terminus of the TRP domain should be at least included as the necessary minimal PIP_2_-dependent gating pathway for the specific temperature threshold range and sensitivity and the reasonable systematic thermal instability.

### 2.3 Preparation of the systematic fluidic noncovalent interaction mesh network maps

For the thermoring structural analyses of a systematic fluidic grid-like mesh network of mTRPM8 along the PIP_2_-dependent minimal gating pathway of one subunit, all the potential stereo-selective or regio-selective inter-domain diagonal and intra-domain lateral noncovalent interactions such as salt/metal bridges and π interactions and H-bonds, once filtered as described previously (Tables S1-3), were geometrically mapped as grids. ^24-28^ By using the Floyd-Warshall algorithm, the grid size was constrained as the minimal number of the total side chains of free residues in protein or atoms in the bound lipid, which did not engage in any noncovalent interaction in a grid. ^29^ In this way, both the shortest direct and reverse paths were obtained to link two ends of a noncovalent interaction. For example in Fig.1A, based on a grid-like biochemical reaction mesh network, a direct path length from F874 to Y908 was zero owing to a π−π interaction between them. However, another shortest reverse path directed from Y908 to F881, W877 and back to F874 via three π interactions. Because no free residue participated in any noncovalent interaction during this cycle, the grid size was zero. Once each noncovalent interaction was tracked by a constrained grid size and the unshared sizes were marked in black, a grid with an x-residue or atom size was denoted as Grid_x_, and then all the noncovalent interactions and grid sizes along the PIP_2_-dependent minimal gating pathway of one subunit were summed in black and cyan circles beside the mesh network map, respectively, for the computation of the melting temperature threshold (T_m_), the systematic thermal instability (T_i_) and the defined structural thermo-sensitivity (Ω_10_).

**Figure 1.**
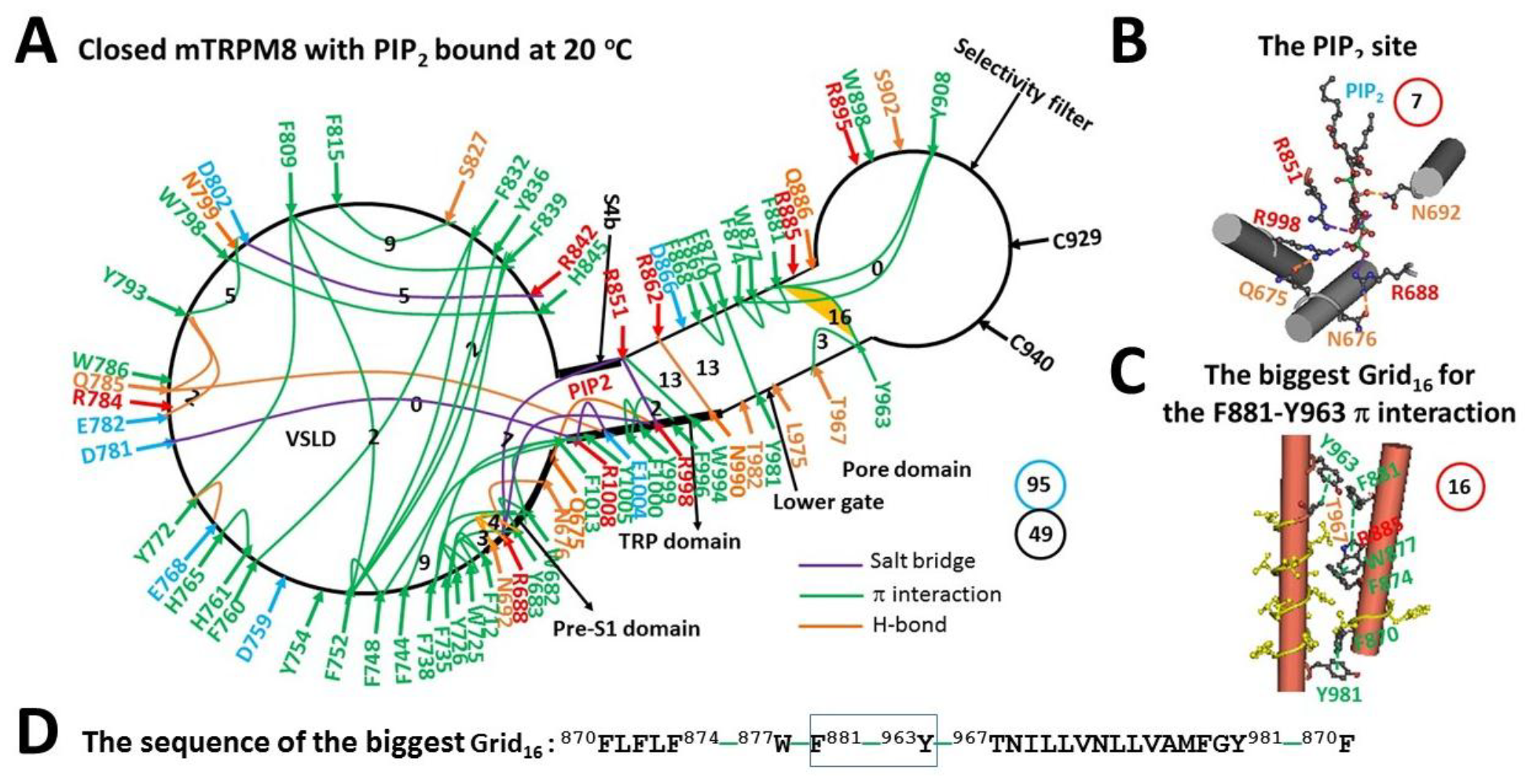
The grid-like noncovalently interacting mesh network along the PIP_2_-dependent minimal gating pathway of closed mTRPM8 with PIP_2_ bound at 20 °C. A, The topological grids in the systemic fluidic grid-like mesh network. The cryo-EM structure of one subunit in homotetrameric closed mTRPM8 with PIP_2_ bound at 20 °C (PDB ID, 8E4N) was used for the model. The pore domain, the selctivity filter, S4b, the TRP domain, the VSLD and the pre-S1 domain are indicated in black. Salt bridges, π interactions, and H-bonds between pairing amino acid side chains along the PIP_2_-dependent minimal gating pathway from Q675 to F1013 are marked in purple, green, and orange, respectively. The grid sizes required to control the relevant noncovalent interactions were constrained with graph theory and labeled in black. The F881-Y963 bridge in the biggest Grid_16_ was highlighted. The calculated total grid sizes and noncovalent interactions are shown in the cyan and black circles, respectively. B, The structure of the PIP_2_ pocket. C, The structure of the biggest Grid_16_ in the S5/S6 interface. D, The sequence of the biggest Grid_16_ to control the F881-Y963 π interaction in the blue box.

### 2.4 Equations

When a 20-base loop and two G-C base pairs in the stem form a DNA hairpin thermo-sensor, an initial T_m_ for loop unfolding is 34 °C, and then increased by 10 °C with a more G-C base pair or five more bases included. ^23^ Similarly, for the thermorings to undergo rate-limiting thermal unfolding from the biggest grid to the smallest one along the PIP_2_-dependent minimal gating pathway, the T_m_ of the given grid could be in turn calculated by using the following equation as described and examined previously: ^24-28^

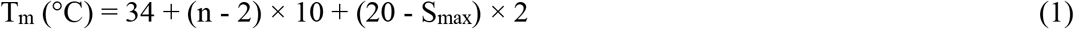

where, n is the total number of the grid size-controlled basic H-bonds (1 kcal/mol each) equivalent to the noncovalent interactions in a given grid, and S_max_ is the size of the given grid. Clearly, a smaller grid size or more equivalent H-bonds are necessary for higher heat capacity in a grid.

Based on the above equation, the more G-C base pairs in the stem or the shorter poly-A loop, the higher T_m_ of the DNA hairpin. ^23^ Thus, in either gating state, the systematic thermal instability (T_i_) along the PIP_2_-dependent minimal gating pathway was reasonably defined using the following equation as described and examined previously: ^24-28^

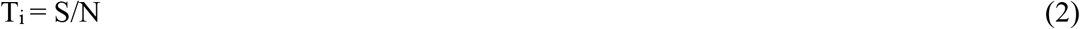

where, along the same PIP_2_-dependent minimal gating pathway of one subunit in a given gating state, S is the total grid sizes and N is the total noncovalent interactions. In this way, the lower the T_i_, the less the compact conformational entropy in the system.

In a temperature range ΔT, for an enthalpy-driven closed-to-open transition, assuming the chemical potential of a grid as the maximal potential for equivalent residues in the grid to form the tightest β-hairpin with the smallest loop via paired noncovalent interactions, ^30^ the grid-based structural thermo-sensitivity (Ω_ΔT_) of a single ion channel can be defined and computed using the following equations:

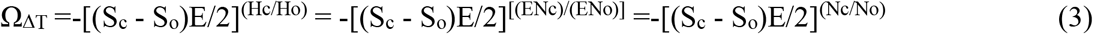

where, along the same PIP_2_-dependent minimal gating pathway of one subunit, N_c_ and N_o_ are the total noncovalent interactions, S_c_ and S_o_ are the total grid sizes, and H_c_ and H_o_ are the total enthalpy included in noncovalent interactions in the closed and open states, respectively. E is the energy strength of a noncovalent interaction in a range of 0.5-3 kcal/mol. Usually, E is 1 kcal/mol. ^31^ Thus, Ω_ΔT_ actually mirrors a thermo-induced change in the total chemical potential of grids upon a thermo-evoked change in the total enthalpy included in noncovalent interactions from a closed state to an open one along the same PIP_2_-dependent minimal gating pathway of one subunit.

When ΔT=10 °C, Ω_10_ could be comparable to the functional thermo-sensitivity (Q_10_) of a single ion channel. Q_10_ during thermal activation was computed using the following equation:

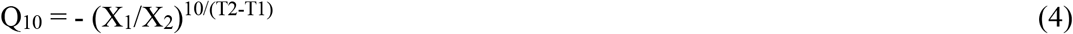

where, X_1_ and X_2_ are open probability (P_o_) values or reaction rates of mTRPM8 at temperatures T1 and T2 (measured in kelvin), respectively.

## 3. RESULTS

### 3.1 Identification of a PIP_2_-dependent cooling switch at 32-37 °C in mTRPM8

For a given PIP_2_-dependent minimal gating pathway from Q675 in the pre-S1 domain to F1013 in the TRP domain of mTRPM8 at 20 °C, nine H-bonds, thirty-four π interactions and six salt bridges shaped a systematic fluidic grid-like mesh network (Fig. 1A). When the total noncovalent interactions and grids size were 49 and 95, respectively (Fig. 1A, Table S1). the grid-based systematic thermal instability (T_i_) was 1.94 (Table 1).

**Table 1.** New PIP_2_-dependnet parameters of the TRPM8 bio-thermometer based on the grid thermodynamic model. The comparative parameters are highlighted in bold.

| Construct | mTRPM8 |  |  |
| --- | --- | --- | --- |
| PDB ID | 8E4L | 8E4M | 8E4N |
| Ca<sup>2+</sup> | bound | bound | free |
| PIP<sub>2</sub> | bound | bound | bound |
| Sampling temperature, °C | **20** | **20** | **20** |
| Gating state | open | closed | closed |
| Name of the biggest grid | Grid<sub>22</sub> | Grid<sub>21</sub> | Grid<sub>16</sub> |
| Biggest grid size ($S_{max}$) | 22 | 21 | 16 |
| Equivalent basic H-bonds for $S_{max}$ | 1.0-1.5 | 1.0-1.5 | 1.0-1.5 |
| Total noncovalent interactions | 51 | 49 | 49 |
| Total grid sizes, a. a./atoms | 74 | 93 | 95 |
| Systematic thermal instability ($T_i$) | 1.45 | 1.90 | 1.94 |
| Calculated $T_m$, °C | **20-25** | **22-27** | **32-37** |
| Measured threshold $T_{th}$, °C | **23-29** | **23-29** | **31-37** |
| Calculated $\Omega_{10, min}$ at E=0.5 kcal/mol | | -4.46 | |
| Calculated $\Omega_{10, mean}$ at E=1.0 kcal/mol | | **-8.68** | |
| Calculated $\Omega_{10, max}$ at E=3.0 kcal/mol | | -24.9 | |
| Measured $Q_{10}$ | | **-9.16** | |
| Ref. for measured $T_{th}$ and $Q_{10}$ | [5] | [5] | [5] |

Of special note, in the presence of 1 mM PIP_2_, ^22^ PIP_2_ was packaged by R668 and N692 in the pre-S1, R851 at S4b, and R998 in the TRP domain via three salt bridges and an H-bond (Fig. 1B). In this case, the biggest Grid_16_ was outstanding in the S5/S6 interface. It had a 16-residue size via the shortest path from F881 to Y963, T967, Y981, F870, F874, W877 and back to F881 (Fig.1C). When 1.0-1.5 equivalent H-bonds sealed this biggest Grid_16_ to hold the F881-Y963 π interaction in the S5/S6 interface (Fig. 1C), the calculated melting temperature threshold (T_m_) ranged from 32 to 37 °C, which was close to the experimental threshold range of 31-37 °C of mTRPM8 for cold activation in the presence of high concentration of PIP_2_ (>500 μM). ^5^ Thereafter, the biggest Grid_16_ may act as a start cooling switch in TRPM8 to initiate channel activation in an optimal temperature range 32-37 °C. However, it could not be broken at 20 °C for channel opening.^22^

### 3.2 Addition of Ca^2+^ and cooling agent C3 decreased the activation threshold range of mTRPM8 to 22-27 °C

It has been reported that Ca^2+^ binding cannot change the temperature threshold for cold-evoked activation of of mTRPM8.^32^ On the other hand, the C3-evoked TRPM8 activity at room temperature (22-24 °C) is not desensitized by Ca^2+.^. ^22^ However, mTRPM8 is not structurally open in the presence of C3 and Ca^2+^ and PIP_2_ at 20 °C. ^22^ Therefore, it is necessary to test the effect of C3 on the activation threshold of mTRPM8. When Ca^2+^ bridged E782, Q785, N799 and D802 together, the Y793-N799 π interaction and the E782-Y793-Q785 H-bonds were broken (Figs. 1A & 2A). Although C3 was bound against Y745 in the voltage sensor-like domain (VSLD), ^22^ it did not link any two residues together via a noncovalent interaction (Fig. 2A). In contrast, the R851-PIP_2_ salt bridge was replaced with the S850-PIP_2_ H-bond (Fig.2B). As a result, when the total noncovalent interactions and grid sizes were 49 and 93, respectively, the resultant systematic thermal instability (T_i_) decreased from 1.94 to 1.90 (Fig. 2A, Table 1). What is more, the biggest Grid_21_ appeared in the S5/S6 interface to control the F881-T967 H-bond. It had a 21-residue size via the shortest path from F881 to W877, F874, F870, F869, D866, Y981, T967 and back to F881 (Fig 2. C-D). With 1.0-1.5 equivalent H-bonds sealing it, the calculated melting temperature threshold range was 22-27 °C, which was close to the experimental range of 23-29 °C in the presence of 100 μM PIP_2_. ^5^ However, it was still higher than the sampling temperature 20 °C of the structural data, thereby leaving the channel closed. ^22^

**Figure 2.**
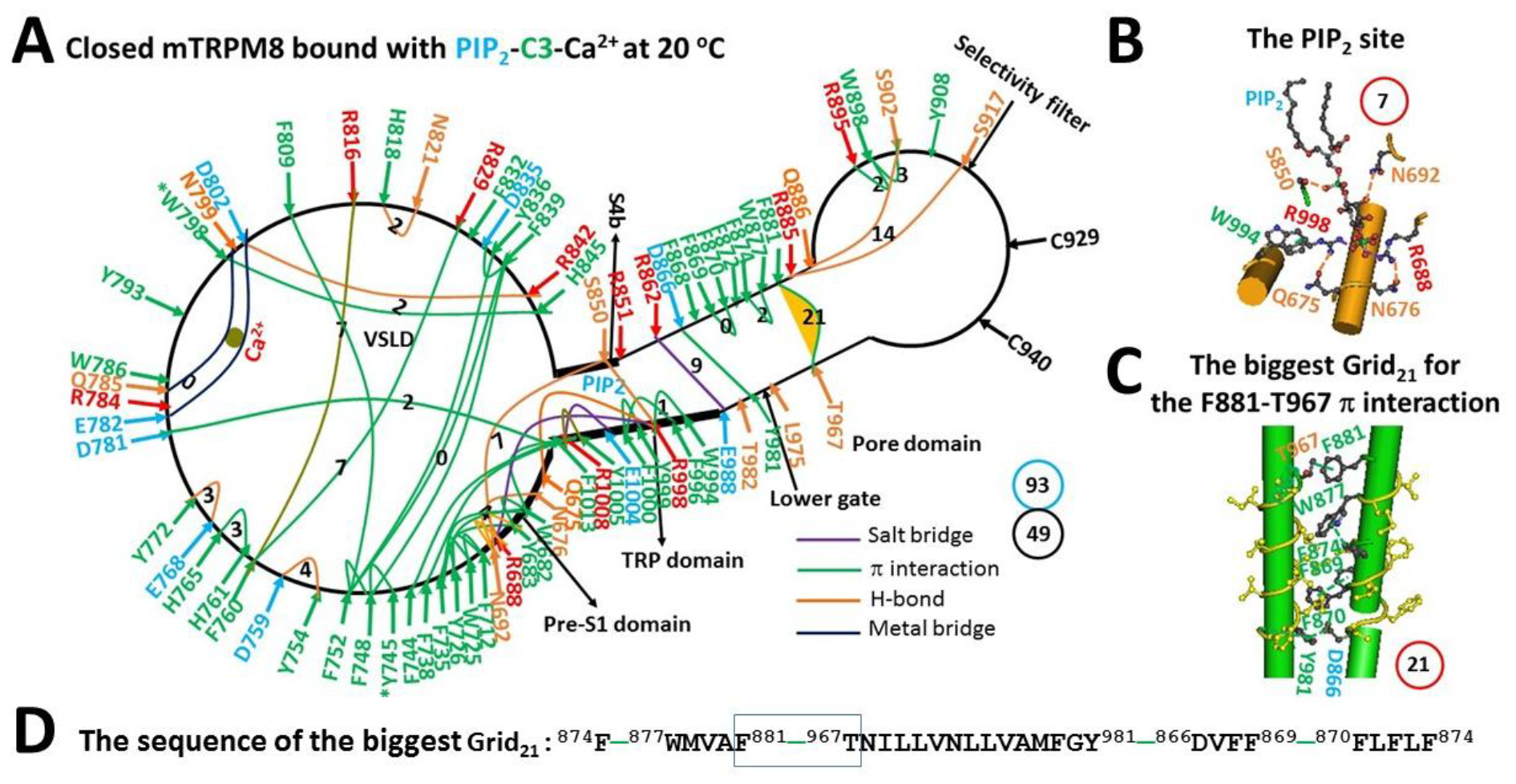
The grid-like noncovalently interacting mesh network along the PIP_2_-dependent minimal gating pathway of closed mTRPM8 with Ca^2+^, PIP_2_ and C3 bound at 20 °C. A, The topological grids in the systemic fluidic grid-like mesh network. The cryo-EM structure of one subunit in homotetrameric closed mTRPM8 with Ca^2+^, PIP_2_ and C3 bound at 20 °C (PDB ID, 8E4M) was used for the model. The pore domain, the selctivity filter, S4b, the TRP domain, the VSLD and the pre-S1 domain are indicated in black. Salt bridges, π interactions, and H-bonds between pairing amino acid side chains along the PIP_2_-dependent minimal gating pathway from Q675 to F1013 are marked in purple, green, and orange, respectively. The grid sizes required to control the relevant noncovalent interactions were constrained with graph theory and labeled in black. The F881-T967 π bridge in the biggest Grid_21_ was highlighted. The calculated total grid sizes and noncovalent interactions are shown in the cyan and black circles, respectively. B, The structure of the PIP_2_ binding site. C, The structure of the biggest Grid_21_ in the S5/S6 interface. D, The sequence of the biggest Grid_21_ to control the F881-T967 π interaction in the blue box.

### 3.3. Further Addition of AITC Opened mTRPM8 at the Matched Threshold 20 °C

In the presence of not only Ca^2+^ and PIP_2_ and C3 but also AITC at 20 °C, mTRPM8 was open with several changes in the systematic fluidic grid-like noncovalent interaction mesh network (Fig. 3A). First, at the Ca^2+^ site, W786-Y793-W798 π interactions and the Y793-E782 and D781-R784 H-bonds were formed. Second, C3 and AITC were bound against Y745 and W798 in the VSLD, respectively. ^22^ However, these two chemical agonists failed to connect any two residues together via a noncovalent interaction (Fig. 3A). Third, PIP_2_ was bound again to R688, R851, and R998, forming a Grid_7_ (Fig. 3B). Fourth, along with the PIP_2_ binding, the D781-F1013 π interaction was disrupted to allow the gating Grid_8_ to be born with an 8-residue size via the shortest path from H845 to R851, W994, R998, Y999, F1000, E1004 and back to H845 (Figs. 3A & 3D). Fifth, when R851 formed a π interaction with W994 and an H-bond with D991 in the TRP domain, the D866-T982 H-bond and the F870-L975 π interaction in the S5/S6 interface were created to form the thermo-stable anchor Grid_7_ in favor of the active lower S6 gate (Fig. 3A). It had a 7-residue size from D866 to F868, F869, F870, L975, T982 and back to D866 (Fig. 3D). Meanwhile, the biggest Grid_22_ in the pore domain had a 22-residue size to stabilize the open state via the shortest path from F874, W877, F881, Y963, R885, Y908 and back to F874 (Figs. 3C-D). When 1.0-1.5 equivalent H-bonds sealed it, the calculated T_m_ ranged from 20 to 25 °C, which matched the sampling temperature 20 °C of the open state. ^22^ As the total noncovalent interactions and grids size were 51 and 74, respectively (Fig. 3A, Table S3), the grid-based systematic thermal instability (T_i_) further decreased to 1.45 (Table 1). For channel opening upon the addition of AITC, the calculated mean Ω_10_ of -8.68 was close to the experimental Q_10_ of -9.16 at 100 μM PIP_2_ (Table 1)._5_

**Figure 3.**
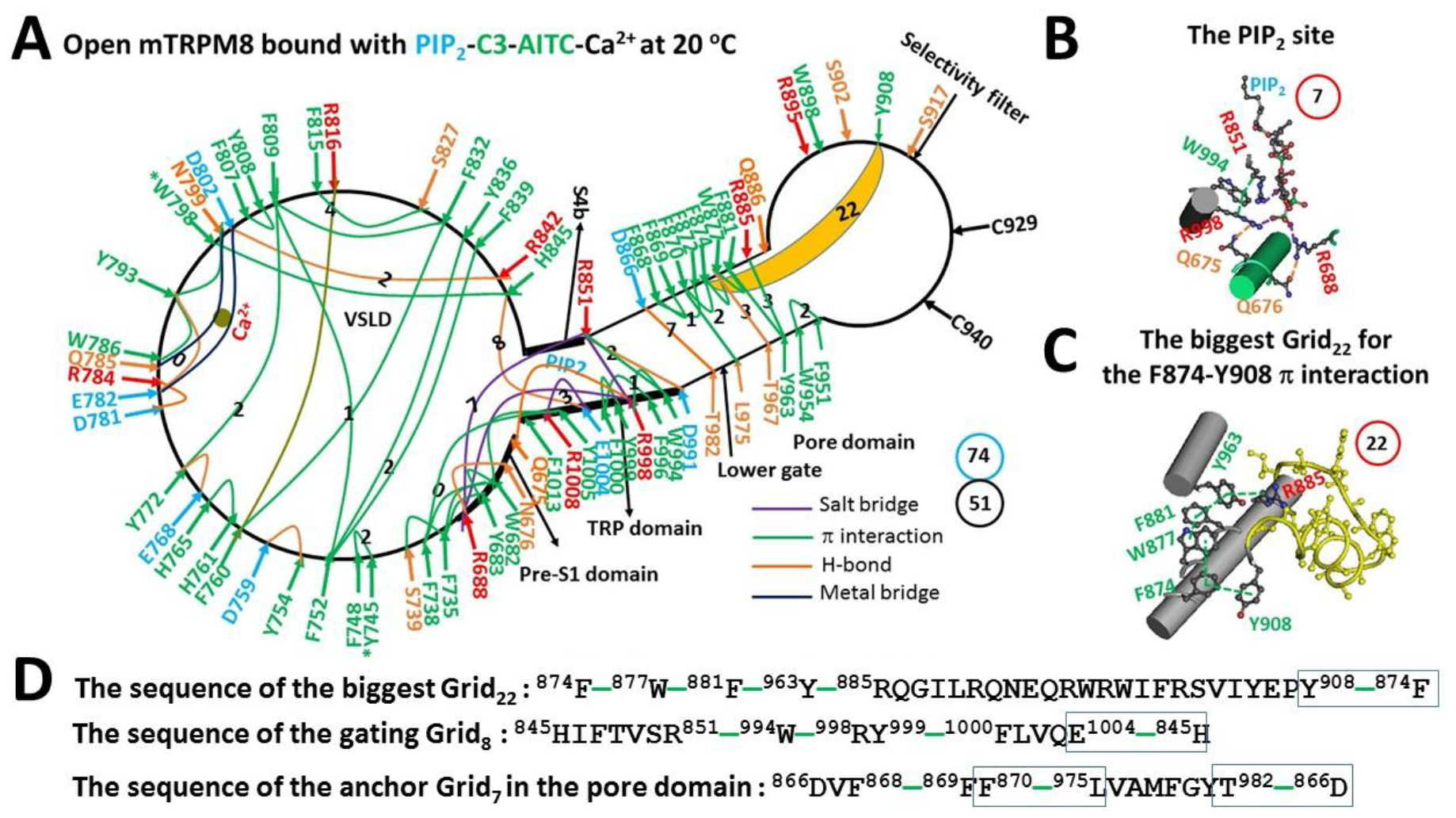
The grid-like noncovalently interacting mesh network along the PIP_2_-dependent minimal gating pathway of open mTRPM8 with Ca^2+^, PIP_2_, C3 and AITC bound at 20 °C. A, The topological grids in the systemic fluidic grid-like mesh network. The cryo-EM structure of one subunit in homotetrameric open mTRPM8 with Ca^2+^, PIP_2_, C3 and AITC bound at 20 °C (PDB ID, 8E4L) was used for the model. The pore domain, the selctivity filter, S4b, the TRP domain, the VSLD and the pre-S1 domain are indicated in black. Salt bridges, π interactions, and H-bonds between pairing amino acid side chains along the PIP_2_-dependent minimal gating pathway from Q675 to F1013 are marked in purple, green, and orange, respectively. The grid sizes required to control the relevant noncovalent interactions were constrained with graph theory and labeled in black. The F874-Y908 bridge in the biggest Grid_22_ at 20 °C was highlighted. The calculated total grid sizes and noncovalent interactions are shown in the cyan and black circles, respectively. B, The structure of the PIP_2_ binding site. C, The structure of the biggest Grid_22_ in the pore domain. D, The sequence of the biggest Grid_22_ to control the F874-Y908 π interaction in the blue box; the sequence of the gating Grid_8_ to control the H845-E1004 H-bond in the blue box; and the sequence of the anchor Grid_7_ in the pore domain to control the D866-T982 H-bond and the F870-L975 π interaction in the blue boxes.

## 4. DISCUSSION

The release of the lipid phosphatidylcholine (PC) or phosphatidylinositol (PI) from the vanilloid site of TRPV3 or TRPV1 is required for heat activation, respectively.^33-34^ In contrast, the cold ativation of TRPM8 needs the lipid PIP_2_. ^5-6, 9, 22^ Although the role of PC or PI in heat response and sensitivity of TRPV3 or TRPV1 has been illuminated recently, ^27-28^ much less is known about how the interaction of TRPM8 with PIP_2_ governs the specific temperature threshold and sensitivity for cold sensing. In this computational study, three PIP_2_-dependent biggest grids in or near the S5/S6 interface of mTRPM8 were identified in the presence of the high concentration of 1 mM PIP_2_. Futher studies showed that the different interactions of mTRPM8 with PIP_2_ in response to the addition of Ca^2+^ and the chemical agents C3 and AITC created declined melting temperature threshold ranges, which were calculated from those PIP_2_-dependent biggest grids. When two threshold ranges in both open and pre-open closed states were similar, the channel could be opened with the matched thermosensitivity and the decrease in the systemeratic thermal instability. Finally, the intact anchor near the lower gate was necessary for channel opening with the active selectivity filter. Therefore, this study may promote our understanding about the role of the lipids in regulating distinct thermorings for specific temperature thresholds and sensitivity in thermo-gated TRP channels.

### 4.1. Interaction of TRPM8 with PIP_2_ Determined the Temperature Threshold Range for Channel Opening

The temperature threshold ranges have been shown to increase from 23-29 °C to 31-37 °C when the concentration of PIP_2_ raised from 0.1 to 0.5 mM. ^5^ However, the underlying structural motif is unknown. This computatinal study identified two biggest grids in the systemic fluidic grid-like noncovalently interacting mesh networks of mTRPM8 in the presence of 1 mM PIP_2_. In the absence of the cooling agent C3, PIP_2_ was found tightly bound with R688 and N692 in the pre-S1 domain, R851 at S4b, and R998 in the TRP domain via three salt bridges or one H-bond (Fig.1B). In this case, the biggest Grid_16_ in the S5/S6 interface generated the calculated melting temperature threshold range from 32 to 37 °C, which was similar to the measured activation threshold range from 31 to 37 °C (Table 1). ^5^ In the presence of the cooling agrent C3, the salt bridge between R851 and PIP_2_ was substituted by the S850-PIP_2_ H-bond (Fig. 2B). In that case, another biggest Grid_21_ in the S5/S6 interface had the calculated melting temperature threshold range from 22 to 27 °C, which was close to the experimental threshold range from 23-29 °C (Table 1). ^5^ Thus, it is possible that PIP_2_ may have a similar interaction with TRPM8 for the biggest Grid_21_ to be a temperature sensor in the pore domain when the concentration declines to 0.1 mM.

On the other hand, the temperature sensation of manmalian TRPM8 is reportedly regulated by the initial or ambient temperature. ^32^ For example, the rise in the exposed temperature from 30 to 40 °C can increase the temperature threshold for the activation of human or rat TRPM8 (hTRPM8 or rTRPM8, respectively) from 28.3 °C to 33.8 °C in *HEK293* cells expressing hTRPM8. ^32^ Further, reducing the level of cellular PIP_2_ or the R1008Q mutation in the TRP domain attenuates or eliminates the ambient temperature-induced changes in the cold temperature threshold. ^32^ Therefore, the interaction of TRPM8 with PIP_2_ plays a pivotal role in modifying the temperature threshold for channel activation. At the ambient temperature 40 °C, the higher level of cellular PIP_2_ may allow the tight interaction in the pre-S1/S4b/TRP interfaces to produce the biggest Grid_16_ or the similar one with a higher temperature threshold 33.8 °C (Fig.1B-D, Table 1). In contrast, at lower pre-determined temperature 30 °C, cellular PIP_2_ may decease the level so that the relatively loose binding to the pre-S1/S4b/TRP interfaces could trigger the formation of another biggest Grid_21_ or the similar one with the lower threshold 28.3 °C (Fig. 2B-D, Table 1). In good agreement with this proposal, in the former case, R1008 in the TRP domain interacted with both D781 in the VSLD and nearby E1004 via the salt bridges (Fig. 1A). However, in the latter case, the D781-R1008 salt bridge was replaced by the D781-F1013 π interaction (Fig. 2A). Thereafter, the R1008Q mutation may allow the loose binding of PIP_2_ to TRPM8 to have the similar lower threshold for channel activation no matter whether the exposed temperature increased from 30 to 40 °C.^32^ Since the cellular PIP_2_ level did decline upon a temperature decrease from 25 to 10 °C, ^35^ it is reasonable that the open state of TRPM8 could be stabilized at the maximal activity temperature 8.2 °C. ^1-2^ In that case, the stimulatory H-bond between H845 in the VSLD and E1004 in the TRP domain may be enough to allow the gating Grid_8_ to hold the open state even at the lower PIP_2_ level once TRPM8 has been activated (Figs. 3A & 3D).

### 4.2 Implication of TRPM8 activation by LPLs and menthol

In addition to PIP_2_, a small dose (3 μM) of long-chain intracellular LPLs can also directly activate TRPM8 even at normal body temperature. This activation has prominent prolongation of channel openings and can be suppressed by protons at pH 6 but not by Mg^2+^ ^10-11^. In this case, PIP_2_ may not be involved so that either or both of free PIP_2_ and AITC sites may hold LPLs with smaller head groups, stablizing the stimulatory interaction between S4b and the TRP domain for channel opening at elevated temperature (Fig. 3B). ^6, 22^

In contrast, menthol may share the same pocket with C3. The binding of a single menthol molecule to R841 on S4 has been proposed to stimulate TRPM8 activity. ^36^ However, that model cannot explain the Y1005-dependent menthol upregulation of TRPM8 with a low efficacy but a high coefficient of a dose response at membrane hyperpolarization, ^19, 37, 38^ and the ligand stereoselectivity of the LNI1009PAA mutant. ^19^, Thus, in addition to H-bonding with R841, an H-bonded homochiral menthol dimer with head-to-head or head-to-tail is still required to push Y1005 against the anchors L841 and I844 on S4 of mTRPM8 for those experimental observations. ^20^ In this way, the tighter interaction between PIP_2_ and TRPM8 may be allowed to increase the melting temperature threshold for channel opening. Further structural data are necessary to reveal the LPLs or menthol sites and their roles in stimulating TRPM8 opening at higher temperature.

### 4.3 Overlapped Temperature Thresholds between Open and Pre-Open Closed States Were Required for Cold-Sensing

The open state of mTRPM8 in the presence of high concentration of PIP_2_ at 20 °C had the biggest Grid_22_ for the melting temperature threshold range of 20-25 °C (Fig. 1, Table 1). Thereby, even if the channel can be opened at 32-37 °C in the presence of 0.5 mM PIP_2_, ^5^ the biggest Grid_22_ would have been melt at 32-37 °C and hence another biggest grid may be necessary to match the higher T_m_ of 32-37 °C for channel opening. In contrast, the pre-open closed state of mTRPM8 with Ca^2+^ and C3 bound in the presence of 1 mM PIP_2_ had the biggest Grid_21_ for the calculated T_m_ range from 22 to 27 °C (Fig. 2C-D). Therefore, the biggest Grid_22_ in the open state could adapt a temperature range from 22 to 25 °C even if AITC was absent (Fig. 4). In other words, the overlapped thresholds between open and pre-open closed states may be necessary for cold activation of TRPM8. Given that the threshold is 25 °C at the cellular PIP_2_ concentration of 10 μM, ^5,39^ further structural investigations of closed and open mTRPM8 at 10 μM PIP_2_ around 25 °C may be required for the detailed understanding about how the TRPM8 bio-thermometer senses cold under the native physiological condition.

**Figure 4.**
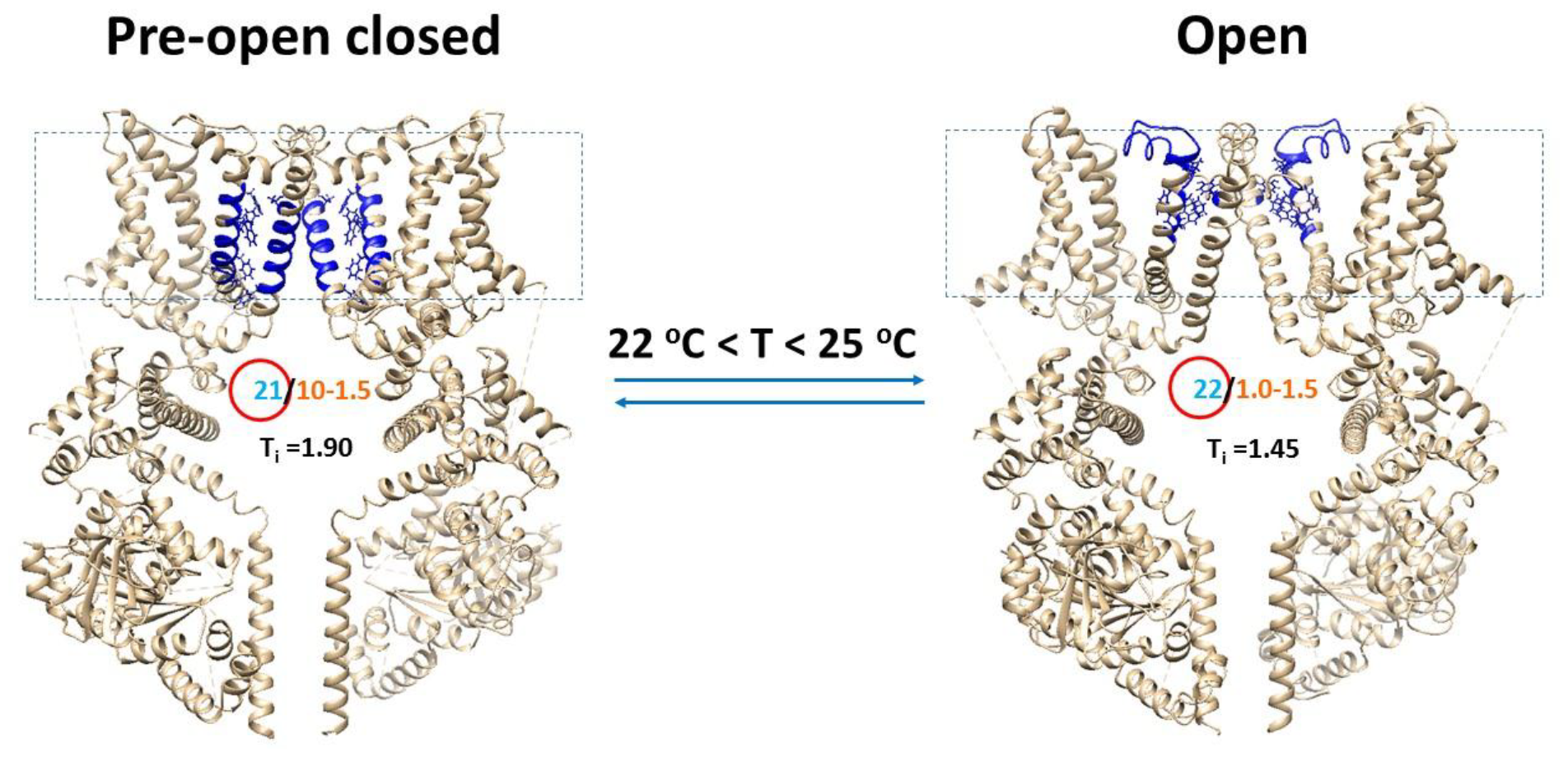
The tentative cold switches in the TRPM8 biothermometer. A, The homo-tetrameric cryo-EM structure of mTRPM8 with (PDB ID, 8E4L) or without (PDB ID, 8E4M) AITC bound in the presence of Ca^2+^ and PIP_2_ and C3 at 20 °C was used for the model of the open or pre-open closed state, respectively. For a convenient view, only two opposite subunits are shown. The dashed rectangle is the membrane area. The PIP_2_-dependent biggest Grid_21_ in the pre-open closed state and the biggest Grid_22_ in the open state have the overlapped T_m_ range of 22-25 °C for channel opening with the matched temperature threshold range and sensitivity (Table 1).

### 4.4 Thermostable F874-Y908 π interaction in the pore domain was needed for the active selectivity filter

Previous chimera study of TRPM8 in cold-sensitive mouse or rat or African elephant versus cold-insensitive squirrel or hamster or chicken or emperor penguin showed that the outer pore loop is thermosensitive. ^40-42^ Of special note, the R897E mutation in mTRPM8 inhibits the cold response but does not alter the menthol response. ^41^ Further, the Y908A mutation rather than the Y908W/F mutation in the P-helix suppresses the cold-sensing although their response to icilin or menthol is not affected. ^43^ Because Y908 is close to the selectivity filter,^22^ this residue is crucial for the cold-evoked activity. This study indicated that the F874-Y908 π interaction was higly conserved in both closed and open states (Figs. 1A and 3A. Tables S1 and S3). In this regard, such a π interaction may be required to secure the active selectivity filter although the segment from Y925 to P950 of the outer pore loop was unstructured in both closed and open states in the absence of the C829-C940 disulfide bond (Figs. 1A-3A).^22,44^

### 4.5 Thermostable anchor Grid_7_ in the S5-S6 interface was necessary for the active lower S6 gate

D866, T982, F870, L975 are highly conserved in thermosensitive TRPM2-5 and TRPV1-4. ^22,33^ The equivalent T680A mutation (T982 in mTRPM8) has been reported to suppress heat activation of TRPV3. ^45^ Moreover, both equivalent D576-T680 H-bond and F590-L673 π interaction in rat TRPV1 (rTRPV1) or both equivalent D586-T685 H-bond and F580-L678 π interaction in mouse TRPV3 (mTRPV3) are always highly conserved in a thermo-stable anchor grid in both closed and open states. ^27-28^ On the other hand, L975 is close to the lower S6 gate of mTRPM8. ^22, 44^ Accordingly, the D866-T982 H-bond and the F870-L975 π interaction in the anchor Grid_7_ in the S5/S6 interface may be required to secure the active lower S6 gate of mTRPM8 (Figs. 3A & 3D). In consonance with this proposal, the Y981F mutation near T982 in rTRPM8 is not functional but the Y981K/E mutation destabilizes the lower gate, rendering rTRPM8 constitutively active. ^46^ Since these two noncovalent interactions only appeared in the open state of mTRPM8 (Figs. 3A & 3D), the gating rearrangement at the temperature threshold may be necessary to keep the intact anchor Grid_7_ in the S5/S6 interface for channel opening with the active selectivity filter.

### 4.6 TRPM8 opened with the matched thermosensitivity and the decrease in the systematic thermal instability

Although the cold sensitivity of TRPM8 in response to a native PIP_2_ environment is high (Q_10_, 24-35), ^10,21,32^ the calculated structural thermosensitivity for channel opening from the pre-open closed state upon the addition of the chemical agent AITC was still lower in the presence of 1 mM PIP_2_ (Ω_10_, -8.68) (Table 1). As this valuse was similar to the experimental Q_10_ of -9.16 at 0.1 mM PIP_2_ or the temperature coefficient (Q_10_, 9) of the rundown rate of cold response upon excision of the membrane patch into the PIP_2_-free bath solution, ^5, 6^ the physiological PIP_2_ concentration for the cold detection of TRPM8 may be dynamic. ^6,9^ On the other hand, although simple 2-state model or an allosteric model has been proposed for single-channel gating mechanisms of TRPM8 activation by cold and membrane depolarization, ^13, 21^ a more complex 7-state model may be needed for the high temperature sensitivity of TRPM8, ^47^ which cannot be figured out by two closed states and one open state in this study. In fact, in addition to the VSLD, ^40^ the side-chain hydrophobicity in the pore domain has been shown to play a critical role in the thermosensitivity of TRPM8 in different vertebrates. ^42^ However, the segment from Y925 to P950 of the outer pore loop in these three states were mostly unstructured (Figs.1A, 2A & 3A). Hence, the alteration in heat capacity as a result of the changes in not only the total grid sizes along the PIP_2_-dependent minimal gating pathway but also the side-chain hydrophobicity in the pore domain along with multiple gating transitions may be required for such high temperature sensitivity. In addition, recent behavioral studies have shown that phosphoinositide interacting regulator of TRPs (PIRT) can bind to PIP_2_ and is required for normal TRPM8-mediated cold-sensing. ^48^ Therefore, further structural and functional examinations of the PIP_2_-dependent thermosensitivity with or without PIRT bound along with the hydrophobic or hydrophilic change in the pore domain are expected under the physiological condition.

In any way, it is interesting that the similar decrease in the systematic thermal instability (T_i,_ from 1.90 to 1.45) was observed upon cold-evoked TRPM8 opening just as heat-evoked TRPV1 opening (T_i,_ from 1.91 to 1.65) or TRPV3 opening (T_i,_ from 1.88 to 1.20). ^27, 28^ In that regard, the thermo-gated TRP channel opening may be driven by the minimization of the systematic thermal instability. Further cross-examinations of other thermo-sensitive TRP channels may be required.

## 5. CLOSING REMARK

The TRPM8 channel can be activated by either cold or chemical agents such as C3 and AITC. This computational study demonstrated that both physical and chemical stimuli may share the similar gating pathway. Although Ca^2+^ binding was not necessary, PIP_2_ was required to induce the stimulatory interaction at the pre-S1 /voltage sensor/TRP domain interfaces. Further, the overlapped thresholds between open and pre-open closed states may favor the cold-evoked channel activation. Finally, the intact thermal anchor grid near the lower gate was required to secure the active selectivity filter for channel opening. Accordingly, this study may provide important clues as to how the TRPM8 biothermometer uses the PIP_2_-dependent thermorings from the biggest grids to the smallest ones to achieve innocuous-to-noxious cold detection with specific matched temperature thresholds and sensitivity. This study further confirmed the existence of a lipid-dependent temperature sensor in thermo-gated TRP channels. The proposed mechanisms and thermoring concept in this study may facilatate our understanding of activation of TRPM8 by lysophospholipids and menthol.

## Conventions and Abbreviations

AITC: allyl isothiocyanate
C3: cryosim-3 or 1-diisopropylphosphorylnonane
cryo-EM: cryo-electron microscopy
Q_10_: functional thermo sensitivity
LPLs: lysophospholipids
T_m_: melting temperature threshold
P_o_: open probability
Ω_10_: structural thermo-sensitivity
PC: phosphatidylcholine
PI: phosphatidylinositol
PIP_2_: phosphatidylinositol-4,5-bisphosphate
PIRT: phosphoinositide interacting regulator of TRPs
T_i_: systematic thermal instability
TRP: transient receptor potential
TRPM8: TRP melastatin-8
cTRPM8: chicken TRPM8
hTRPM8: human TRPM8
mTRPM8: mouse TRPM8
rTRPM8: rat TRPM8
mTRPV3: mouse TRPV3
rTRPV1: rat TRPV1
VSLD: voltage-sensor-like domain
WS-12: N-(4-methoxyphenyl)-p-menthone-3-carboxamide

## Acknowledgements

The author’s own studies cited in this article were supported by the American Heart Association Grant (10SDG4120011 to GW), National Institute of Health Research Grant 2R56DK056796-10, and Cystic Fibrosis Foundation grant (DAWSON0210).

## Author contributions

G. W. wrote the main manuscript text, prepared table 1 and figures 1-4 and tables S1-S3, and reviewed the whole manuscript and the Supporting Information.

## Data Availability Section

All data generated or analysed during this study are included in this published article.

