## Supplementary Tables S1, S2 and S3 for "PIP_2_-Dependent Thermoring Basis for Cold-Sensing of the TRPM8 Biothermometer"

Running title: Digit codes for cold-gated TRPM8

Guangyu Wang 1, 2*

[1]Department of Physiology and Membrane Biology, University of California School of Medicine, Davis, CA 95616, USA

[2]Department of Drug Research and Development, Institute of Biophysical Medico-chemistry, Reno, NV 89523, USA

**The SI includes:**

**Tables S1, S2, and S3**

**Table S1 Noncovalent interactions along the $PIP_2$-dependent minimal gating pathway from Q675 to F1013 in each subunit of closed mTRPV8 with $PIP_2$ bound at 20 °C (PDB ID, 8E4N)**

| Noncovalent interaction | Cut-off distance | Linked residues |
|---|---|---|
| Salt bridge | 3.2-4 Å | R688-$PIP_2$-R851-$PIP_2$-**R998-$PIP_2$-R668,** D781-R1008, D802-R842, **E1004-R1008** |
| H-bond | <3.9 Å | **Q675-R998, N676-R668,** Y683-N692-R688, E768-Y772, E782-Y793-Q785, Q785-E1004, R862-N990 |
| π–π interaction | 2.65–6.5 Å | **W682-Y683-F735, W682-F735, W682-F738, W682-Y1005,** F712-W725, F712-Y726, W725-Y726, **F738-Y1005,** F744-F1013, F748-F752, F748-F839, **F752-F809,** F752-F832, **F752-Y836, F752-F839,** F760-F832, **H761-H765, Y772-F809, W798-H845, F809-F839,** F868-F869, F870-Y981, F874-W877, F874-Y908, W877-F881, F881-Y963, **F996-F1000, Y999-F1000** |
| cation–π interaction | <6.0 Å | |
| $CH_3$/CH–π interaction | 2.65-3.01 Å | R851-W994, M878-Y908, Y963-T967 |
| Lone pair–π interaction | 3-3.7 Å | Y793-N799, F815-S827 |

Note: Bold interactions were conserved in both closed and open states.

**Table S2 Noncovalent interactions along the $PIP_2$-dependent minimal gating pathway from Q675 to F1013 in each subunit of closed mTRPM8 with $Ca^{2+}$ and $PIP_2$ and C3 bound at 20 °C (PDB ID, 8E4M)**

| **Noncovalent interaction** | **Cut-off distance** | **Linked residues** |
|---|---|---|
| Salt bridge | 3.2-4 Å | **R688-$PIP_2$-R998, E782-$Ca^{2+}$-D802, Q785-$Ca^{2+}$-N799,** R862-E988, **E1004-R1008** |
| H-bond | | **Q675-R998, N676-R688,** Y683-N692, R688-N692, **Y754-D759, E768-Y772, D802-R842,** H818-N821, R688-$PIP_2$-S850-$PIP_2$-R998, R885-S917, Q886-S902 |
| π–π interaction | 2.65–6.5 Å | **W682-Y683, W682-F735, W682-F738, W682-Y1005, Y683-F735,** F712-W725, F712-Y726, W725-Y726, **F738-Y1005,** F744-F1013, **Y745-F748,** F748-F839, **F752-F809, F752-Y836, F752-F839, H761-H765, W798-H845,** F869-F870, **F874-W877, F996-F1000, Y999-F1000** |
| cation–π interaction | <6.0 Å | **F760-R816,** Y1005-R1008 |
| $CH_3$/CH–π interaction | 2.65-3.01 Å | F748-F752, F760-R829, D781-F1013, D835-F839, D866-Y981, F881-T967, R895-W898-S902, **W994-R998** |
| Lone pair–π interaction | 3-3.7 Å | |

Note: Bold interactions were conserved in both closed and open states.

**Table S3 Noncovalent interactions along the $PIP_2$-dependent minimal gating pathway from Q675 to F1013 in each subunit of open mTRPM8 with $Ca^{2+}$ and $PIP_2$ and cooling agents C3 and AITC bound at 20 °C (PDB ID, 8E4L)**

| Noncovalent interaction | Cut-off distance | Linked residues |
|---|---|---|
| Salt bridge | 3.2-4 Å | R688-$PIP_2$-R851-$PIP_2$-**R998-$PIP_2$-R688, E782-$Ca^{2+}$-D802, Q785-$Ca^{2+}$-N799, E1004-R1008** |
| H-bond | <3.9 Å | **Q675-R998, N676-R688, Y754-D759, E768-Y772, D781-R784, E782-Y793, D802-R842,** H845-E1004, R851-D991, D866-T982, W877-T967 |
| π–π interaction | 2.65–6.5 Å | **W682-Y683, W682-F735, W682-F738, W682-Y1005, Y683-F735**, **F738-Y1005, Y745-F748, F752-F809, F752-Y836, F752-F839,** F760-F832, **H761-H765,** Y772-F809, W786-Y793, **W798-H845,** F807-Y808, F809-F832, F868-F869, F868-F872, **F874-W877,** F874-Y908, W877-F881, F881-Y963, F951-W954, **F996-F1000, Y999-F1000** |
| cation–π interaction | <6.0 Å | **F760-R816** |
| $CH_3$/CH–π interaction | 2.65-3.01 Å | Y793-N799, F815-S827, R851-W994, F870-L975, F870-F874, **W994-R998** |
| Lone pair–π interaction | 3-3.7 Å | F735-S739 |

Note: Bold interactions were conserved in both closed and open states.
